# A literature-based meta-analysis of brain-wide electrophysiological diversity

**DOI:** 10.1101/014720

**Authors:** Shreejoy J. Tripathy, Shawn D. Burton, Matthew Geramita, Richard C. Gerkin, Nathaniel N. Urban

## Abstract

For decades, neurophysiologists have characterized the biophysical properties of a rich diversity of neuron types. However, identifying common features and computational roles shared across neuron types is made more difficult by inconsistent conventions for collecting and reporting biophysical data. Here, we leverage NeuroElectro, a literature-based database of electrophysiological properties (www.neuroelectro.org), to better understand neuronal diversity – both within and across neuron types – and the confounding influences of methodological variability. We show that experimental conditions (e.g., electrode types, recording temperatures, or animal age) can explain a substantial degree of the literature-reported biophysical variability observed within a neuron type. Critically, accounting for experimental metadata enables massive cross-study data normalization and reveals that electrophysiological data are far more reproducible across labs than previously appreciated. Using this normalized dataset, we find that neuron types throughout the brain cluster by biophysical properties into 6-9 super-classes. These classes include intuitive clusters, such as fast-spiking basket cells, as well as previously unrecognized clusters, including a novel class of cortical and olfactory bulb interneurons that exhibit persistent activity at theta-band frequencies.

## Introduction

Neurophysiologists have recorded and published vast amounts of quantitative data about the biophysical properties of neuron types across many years of study. Compared to other fields, however, little progress has been made in compiling and cross-analyzing these data, let alone collecting or depositing measurements or raw data 1, 2. It is thus difficult, for example, to determine whether a cerebellar Purkinje cell is more similar to a hippocampal CA1 pyramidal cell or a cortical basket cell without first re-collecting such data in a dedicated experiment, even though thousands of recordings have been made from these neuron types across many laboratories. By analogy to genetics, imagine if genes needed to be re-sequenced every time an investigator wanted to examine genetic homology 3, 4.

The fundamental challenge in comparing electrophysiological data collected across laboratories is twofold. First, unlike genetic sequences 3, 4 or neuron morphologies [5], electrophysiological data is not compiled centrally but remains scattered throughout the vast literature 1, 2. Second, and perhaps more critically, electrophysiological data is collected and reported using inconsistent methodologies and nomenclatures [6]. Thus, if two labs report phenotypic differences for the same neuron type, do these differences reflect true biological differences? Or are they merely the result of methodological differences? These challenges stand as a major barrier to comparison and generalization of results across neuron types, and routinely lead to unnecessary replication of experiments and the slowing of progress 1, 2.

Here, we present a novel approach for integrating and normalizing arbitrarily large amounts of brain-wide electrophysiological data collected across different labs. In contrast to costly ongoing efforts by large institutes to record such data anew 7, 8, 9], our methods capitalize on the immense wealth of data on neuronal biophysics that has already been painstakingly recorded and published. By leveraging the methodological variability inherent in how different labs collect and report biophysical data, we develop statistical methods to disentangle the confounding role of methodological inconsistencies from true biophysical differences among neuron types. This allows us to normalize and compare data collected across labs, including our own, and assess whether neuron types in disparate regions of the brain share common electrophysiological profiles, and thereby fulfill common computational and circuit functions.

This work is of relevance to the broad community of neurophysiologists and computational modelers as it makes large amounts of valuable electrophysiological data easily accessible for subsequent comparison, reuse, and reanalysis. More generally, our work offers a partial solution to the perceived reproducibility crisis in science 10, 11, 12], by demonstrating how data collected using methodologically inconsistent sources can be combined and leveraged to generate novel insights.

## Results

### Building an electrophysiological database by mining the research literature

To make use of the formidable amount of neuronal electrophysiological data present within the research literature, we developed methods to attempt to “mine” such data from the text of published papers. While forgoing the difficulties of recording anew from multiple neuron types and brain areas, such a data-mining approach is not without its own challenges. These challenges include inconsistencies in published neuron naming schemes [6], in how electrophysiological properties are defined and calculated, and in experimental conditions themselves [9]. However, we reasoned that these limitations could potentially be overcome, by capitalizing on the redundancy of published values and the presence of informal community-based reporting standards 13, 6. Our hope was thus to produce a unified dataset of sufficient quality for use in subsequent meta-analyses, and further, that the dissemination of such a resource would encourage better standardization and consistency of future data collection.

We built a database, NeuroElectro, that links specific neuron types to measurements of biophysical properties reported within published research articles (Fig. 1A). Specifically, from 331 articles, we extracted and manually curated information on basic biophysical properties of 97 neuron types recorded during normotypic (i.e., “control”) conditions (Fig. S1A,B). Briefly, our mining strategy follows a three-stage process (detailed in ref. [14]). First, we developed automated text-mining algorithms 15, 16 to identify and extract content related to biophysical properties and experimental conditions. Our algorithms extracted reported mean biophysical measurement values, reflecting pooled values computed across multiple neurons within a type. Second, we manually curated the mined content, taking care to correctly label misidentified neuron types or electrophysiological properties. To help categorize the neuron types recorded within each article, we used the semi-standardized listing of expert-defined neuron types provided by NeuroLex 17, 18. Finally, we manually standardized the extracted electrophysiological values to a common set of units (e.g., GO to MO) and calculation conventions where possible (Fig. S1C,D). We found the accuracy for data categorization and extraction to be 96% overall during a systematic quality control (QC) audit (see Appendix), which we deemed to be of sufficient quality for further meta-analyses.

A sample of the resulting data is shown in Fig. 1 and the dataset in its entirety can be interactively explored through our web interface at http://neuroelectro.org. The dataset reflects known features of several neuron types; for example, cortical basket cells have narrow action potentials [19] and striatal medium spiny neurons rest at relatively hyperpolarized potentials [20]. In this study, we have focused our meta-analyses on six commonly and reliably reported biophysical properties: resting membrane potential, input resistance, membrane time constant, spike half-width, spike amplitude, and spike threshold (abbreviated as V_*rest*_, R_*input*_, T_*m*_, AP_*hw*_, AP_*amp*_, AP_*thr*_, respectively). Other parameters, such as spike afterhyperpolarization amplitude and time course, are recorded in NeuroElectro, but we chose not to not include them in the following analyses due to questions about the consistency of their reporting in the literature.

**Fig. 1.**
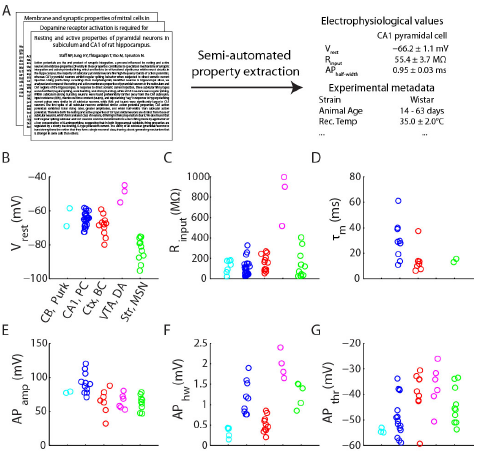
*Schematic of NeuroElectro database construction and example electrophysiological measurements.* A) Semi-automated text-mining algorithms were applied to journal articles to extract neuron type-specific biophysical measurements and experimental conditions. B-G) Example electrophysiological measurements extracted from the research literature for cerebellar Purkinje cells, CA1 pyramidal cells, cortical basket cells, ventral tegmental area dopaminergic cells, and striatal medium spiny neurons (abbreviated as CB, Purk; CA1, PC; Ctx, BC; VTA, DA; and Str, MSN). Each circle denotes the value of the *mean* biophysical measurement value reported within an article.

### Experimental metadata explains cross-study variance among electrophysiological measurements

Our literature-based approach relies on pooling information across articles, which has the inherent advantage of distilling the consensus view of several expert investigators and laboratories. However, data collected under different experimental conditions may not be directly comparable. For example, R_*input*_ tends to decrease as animals age 21, 22. Because NeuroElectro measurements are randomly sampled from the literature, relationships between experimental conditions (“metadata”) and electrophysiological measurements (“data”) should also be reflected within the dataset (Fig. 2A,B). By annotating each electrophysiological measurement in our database with a corresponding set of experimental metadata (Fig. 2A and S2), we were able to address the following three questions. First, can experimental metadata be used to account for and even correct for the variability of data reported across studies? Second, what is the influence of specific experimental conditions (e.g., recording temperature and electrode type) on measurements of biophysical properties? Third, what is the residual variability in reported values for a given neuron type after differences in several experimental conditions have been accounted for?

We used linear regression models to characterize the relationship between electrophysiological measurements and experimental metadata (after appropriate data filtering, like removing data from non-rodent species). We first asked to what extent the variability observed among electrophysiological measurements could be explained by neuron type alone (i.e., how consistent are measurements of the same neuron type from article-to-article). We found that V_*rest*_ was reported fairly consistently (Fig. 2C; adj. R^2^ = 0.6; i.e., 60% of the variability in V_*rest*_ across cells was explained by cell type). However, most properties, such as T_*m*_ and AP_*thr*_, had measurements which differed greatly across studies recording from the same neuron type (adj. R^2^ < 0.25). Thus, there exists a large amount of variance in electrophysiological data that is unexplained by neuron type alone.

We found in many cases, however, that experimental metadata could significantly explain the variability in reported electrophysiological data (Figs. 2D-F, summary in G). For example, knowing whether neurons were recorded using patch versus sharp electrodes explained a substantial fraction of the observed variance in *Ri*_input_, with sharp electrodes yielding on average ∼100 MQ lower *R_input_* than patch electrodes (Fig. 2D). Thus, the dataset inherently reflects a historical controversy when the patch-clamp technique was first introduced and similar large discrepancies were observed in R_*input*_ mea-surements made with patch versus sharp electrodes [23]. Collectively, incorporating multiple experimental metadata factors accounted for considerably more measurement variability than neuron type alone (Fig. 2B; details in Methods). Im-portantly, these regression models provide *quantitative* relationships which can be used as “correction factors” to adjust or *normalize* each electrophysiological measurement for multivariate differences in recording practices across studies (Fig. 2 A,H). Such adjustments are conceptually analogous to “Q10” correction factors, often used to systematically correct for temperature dependent kinetic reaction rates [24].

**Fig. 2.**
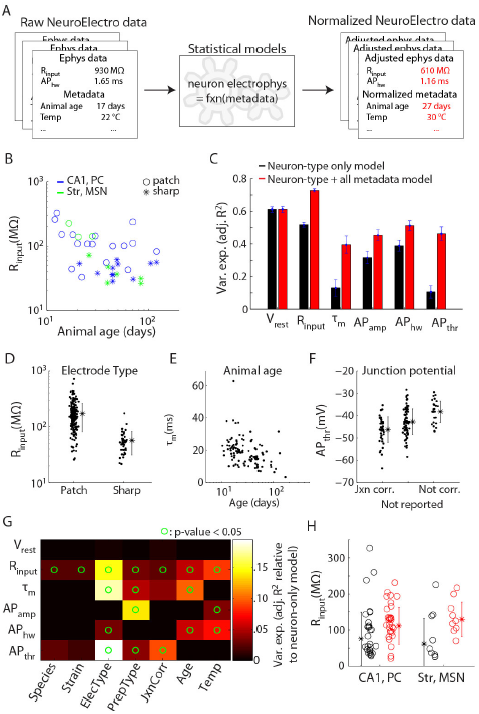
*Methodological differences significantly explains observed variability in literature-mined electrophysiological data.* A) Cartoon illustrating metadata-based NeuroElectro data normalization. B) Example data showing how measured values of *R_input_* vary as a function of recording electrode type and animal age. C) Variance explained by statistical models for each electrophysiological property when only neuron type information is used (black) and when neuron type plus all metadata attributes are used (red). Error bars indicate standard deviations, computed from 90% bootstrap resamplings of the entire dataset. D-F) Example relationships between specific metadata predictors and variation in electrophysiological properties. Dots show model-adjusted electrophysiological measurements after accounting for specific differences across neuron types. Panel F refers to correction of liquid junction potential (“jxn”). Asterisks indicate population mean and error bars indicate s.d. G) Influence of individual metadata predictors in helping explain variance in specific electrophysiological properties. Heatmap values indicate relative improvement over the model that includes neuron type information only. Circles indicate where the regression model including a metadata attribute was statistically more predictive than the model with neuron type information alone (p < 0.05; ANOVA). H) Example data before (black) and after using statistical models to adjust for differences in metadata among electrophysiological measurements (red). Measurements become less variable and skewed after adjustment for methodological differences.

As a caveat, we note that there still remained a considerable amount of unexplained variance in electrophysiological measurements, even after metadata adjustment. This variance likely reflects: 1) within-type neuronal variability 25, 26; 2) additional experimental conditions not yet considered, like recording solution contents (e.g., see Fig. S3) or overall preparation and recording quality; and 3) subtle differences in how investigators define electrophysiological properties (e.g., see Fig. S4 and Discussion).

**Fig. 3.**
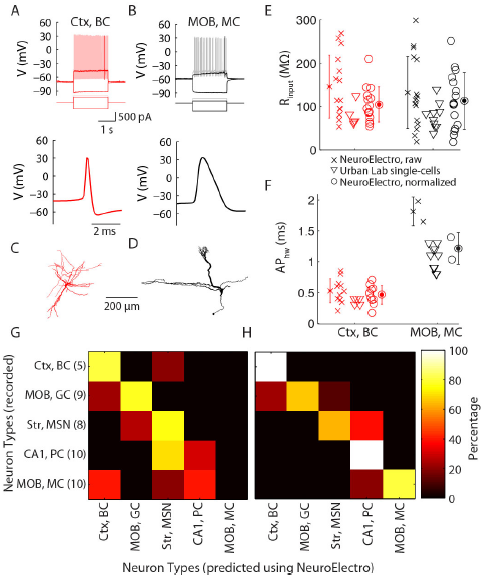
*Direct comparison of NeuroElectro measurements to* de novo *recordings.* A,B) Representative recordings of a neocortical basket cell (A) and a main olfactory bulb mitral cell (B), showing responses to hyperpolarizing, rheobase, and suprathreshold current injections (top), and AP waveform (bottom). C,D) Morphologies for cells in A and B. E, F) Database measurements for mitral and basket cells before (crosses) and after (circles) metadata normalization and cor-responding Urban Lab single-cell measurements (triangles). Error bars indicate s.d., computed across database measurements within a neuron type. G,H) Confusion matrices highlighting classification of each recorded single-cell to corresponding aggregate NeuroElectro neuron type for the raw (G) or metadata-normalized (H) NeuroElectro dataset. Matrix y-axis indicates recorded neuron identity and number within parentheses indicates n of recorded single-cells per neuron type. X-axis indicates the predicted neuron type based on biophysical similarity to NeuroElectro (i.e., perfect classification is a diagonal along matrix).

### Experimental validation of NeuroElectro data before and after metadata normalization

To directly validate NeuroElectro dataset measurements and our metadata normalization procedure, we recorded from a subset of commonly studied neuron types, including CA1 pyramidal cells, main olfactory bulb mitral cells and granule cells, neocortical basket cells, and striatal medium spiny neurons (Fig. 3A-F and S5). Critically, to compare our *de novo* recordings to NeuroElectro, we needed to first statistically normalize the NeuroElectro measurements to the experimental conditions used in our lab - namely, our use of whole-cell patch-clamp recordings near physiological temperatures in acute slices from young-adult mice.

Comparing our single-cell biophysical measurements to NeuroElectro, we found close agreement to the global mean and variance defining each NeuroElectro neuron type following metadata adjustment (Fig. 3E,F). To quantify this agreement, we used a confusion matrix analysis to classify each recorded cell to the corresponding most similar NeuroElectro neuron type. Experimentally recorded neurons were almost always correctly matched to the corresponding NeuroElectro neuron type after metadata normalization (81% of cells; 34 of 42 neurons; chance = 20%; Fig. 3H), but matched considerably less well when using the raw unnormalized NeuroElectro values (48% of cells; 20 of 42 neurons; Fig. 3G).

We thus conclude that the electrophysiological data and metadata populating NeuroElectro are sufficiently accurate and that individual labs can reasonably expect their own recordings to match NeuroElectro after adjusting for differences in experimental conditions. Moreover, this analysis underscores that: 1) single neurons, even of the same canonical type, are biophysically heterogeneous 25, 26; and 2) incorrectly matched neurons may provide clues to functional similarities across different neuron types.

### Investigating brain-wide correlations among biophysical properties

We next performed a series of analyses on our normalized brain-wide electrophysiology dataset with the goal of learning relationships between biophysical properties and diverse neuron types. To further help reduce the influence of unaccounted-for measurement and methodological variability, we first summarized electrophysiological data at the neuron type level by pooling measurements across articles. Correlating measurements of biophysical properties across neuron types, we observed a number of significant correlations (examples in Fig. 4A,B; summary in C and S6), including correlations expected *a priori,* such as a positive correlation between *R_input_* and *T_*m*_*. We also observed biophysical correlations more difficult to explain via first principles of neural biophysics, such as anti-correlation between R_*input*_ and *AP_amp_* and correlation between AP_*thr*_ and *AP_*hw*_.*

**Fig. 4.**
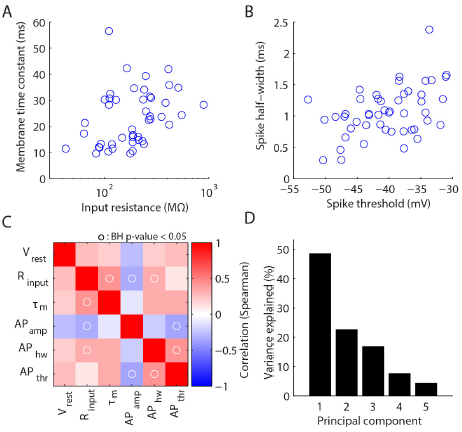
*Exploring correlations between biophysical properties.*A,B) Example data showing pairwise correlations among biophysical properties. Each data point corresponds to measurements from a single neuron type (after averaging observations collected across multiple studies and adjusting for experimental condition differences). C) Correlation matrix of biophysical properties (Spearman's correlation). Circles indicate where correlation of biophysical properties was statistically significant (p!0.05 after Benjamini-Hochberg false discovery rate correction). D) Variance explained across probabilistic principal components of electrophysiological correlation matrix in C.

These correlations led us to use dimensionality reduction techniques to determine if this six parameter description of neuronal diversity could be further simplified. Principal component analysis (using probabilistic PCA, pPCA, to help account for unobserved or “missing” biophysical measurements) showed that 50% of the variance across neuron types could be explained by a single component that largely reflects neuronal size (Fig. 4D and S6B). An additional 22% can be explained by the second PC which roughly reflects basal firing rates and excitability (Fig. S6C,D). This analysis is unique through its focus on *brain-wide neuronal diversity*; moreover, such relationships may differ from previous correlations based on *within neuron type variability* 25, 26.

### Biophysical similarity identifies approximately 6-9 superclasses of neuron types

Lastly, we used NeuroElectro to gain insights into unknown biophysical similarities among diverse neuron types, with the goal of uncovering shared homology of function between different neurons. For example, fast-spiking basket cells populate multiple brain regions yet play similar functional roles within their larger neural circuits 27, 19. Our goal was to use the normalized electrophysiological features to identify additional sets of biophysically similar neuron types which may also share computational functions.

We performed a hierarchical clustering analysis of the neuron types using the metadata-normalized NeuroElectro dataset. Specifically, for each pair of neuron types, we assessed their similarity by comparing the set of six basic biophysical properties defined above. Here, we chose to be agnostic about the relative importance of each biophysical property and weighted them by their relative measurement uncertainty (defined in Fig. 2C). We further mitigated unaccounted-for measurement and methodological variability by focusing on neuron types reported within at least three articles.

Several previously described classes of neuron types emerged from this analysis, validating our unbiased clustering approach (Fig. 5). For example, neocortical and hippocampal basket cells were closely clustered, as were GABAergic medium spiny neurons of both dorsal and ventral striatum. Likewise, we observed distinct clusters of both excitatory neocortical and non-neocortical projection neuron types, differing with respect to their *V_*rest*_.* Further, metadata normalization was critical, as performing the analysis using the unnormalized dataset gave paradoxical results; for example, that CA1 basket cells were more similar to thalamic relay cells than to cortical basket cells (Fig. S7).

**Fig. 5.**
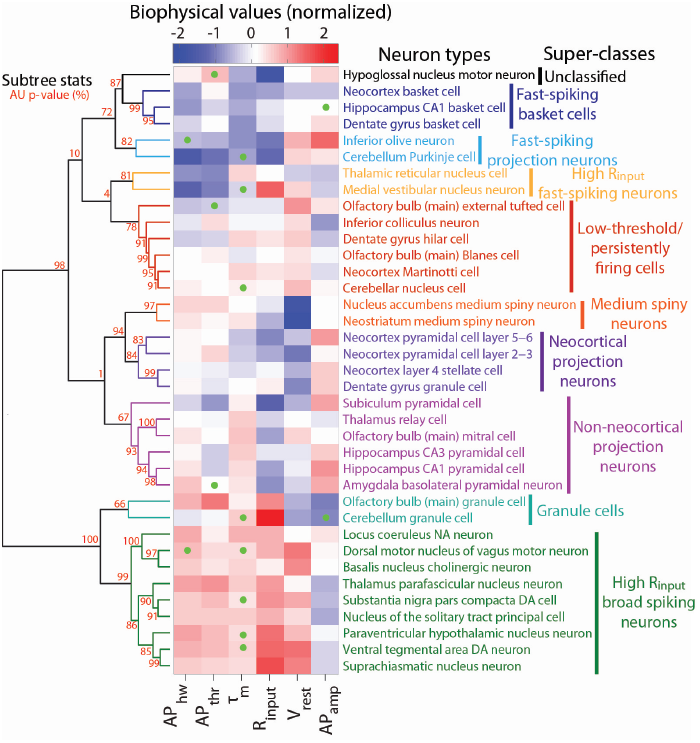
*Hierarchical clustering of diverse neuron types on the basis of biophysical similarity.* Neuron types sorted in order of biophysical similarity (similiarity indicated by levels of dendrogram; dendrogram linkages computed using Ward's method and Euclidean distances). Heatmap values indicate observed neuron type-specific electrophysiological measurements, red (blue) values indicate large (small) values relative to mean across neuron types. Statistical consistency of dendrogram subtrees calculated via bootstrap resampling (red values indicate approximately unbiased (AU) p-values (see Methods); p-values rounded to nearest integer for visualization). Dendrogram subtrees are grouped into neuron type super-classes indicated by text coloring (and are otherwise black) based on p-values and visual inspection. Only neuron types with measurements defined by at least three articles and with at least four (of the six total) biophysical properties reported were used in this analysis. Probabalistic PCA was used to impute unobserved measurements, indicated via green dots on heatmap.

Novel super-classes of neuron types also emerged from our clustering analysis. Foremost, we observed a cluster containing main olfactory bulb Blanes and external tufted cells, dentate gyrus hilar cells, and neocortical Martinotti cells that were defined by a depolarized *V_rest_* and relatively hyperpolarized AP_*thr*_. This is the first report identifying the shared electrophysiological similarity of these neuron types. Intriguingly, each of these neuron types exhibits low-threshold and persistent spiking activity at theta-band frequencies [28, 29, 30, 31], and thus may share the computational function of driving or triggering recurrent network theta rhythms. Similarly, we observed a large cluster of high *R*_*input*_, broad spiking cells from the midbrain and brainstem, including the VTA and locus coeruleus. Though markedly diverse in their combined neurotransmitter phenotype, many of these neuron types nevertheless exhibit similar activity patterns comprised of spontaneous “pacemaker”-like tonic firing 32, 33, a behavior attributable to their distinctively depolarized *V_rest_.*

Across the entire dataset, the major divisions among neuron types tended to be in terms of neuron size and basal excitability (see also Fig. S4E-H). Additionally, we observed a qualitative correspondence between biophysical similarity and gross anatomical position, suggesting that shared precursor lineage may yield similar biophysical properties [34]. While this initial analysis is focused only on simple biophysical properties, the observed “super-classes” are encouraging because they appear to also reflect gestalt spike pattern phenotypes, such as pacemaker or low-threshold firing behaviors. In the future, incorporating additional parameters such as spike AHP amplitudes or ionic currents will likely further refine these super-classes and better define their computational roles 13, 35.

## Discussion

Here, we have developed a general approach for reconciling long-standing methodological inconsistencies that have made brain-wide meta-analyses of electrophysiological data exceedingly difficult. Using semi-automated text-mining [14], we were able to accurately compile considerable amounts of neuronal biophysical data from the vast research literature. In our initial analyses of the extracted data within NeuroElectro, we found that the raw biophysical data values pertaining to the same neuron type were immensely variable across studies. However, the size of this unprecedented collection of electrophysiological data enabled us to explicitly quantify the relationships between experimental conditions and biophysical properties. With these statistical models, we could systematically normalize methodologically-inconsistent data to account for basic differences in experimental protocols thereby revealing the actual features of neuron types.

Following methodological normalization, we obtained, for the first time, a unified reference dataset of neuronal biophysics amenable to brain-wide electrophysiological comparisons. Such metadata normalization was critical for comparing our *de novo* single-cell recordings to NeuroElectro data from other neuron types. Our subsequent meta-analyses uncovered novel electrophysiological correlations and several biophysically-based neuronal super-classes predicted to exhibit similar functionality. For example, we identify a new super-class containing hippocampal, neocortical, and olfactory bulb interneurons capable of persistent theta frequency activity - an emergent behavior attributable, in part, to a uniquely depolarized *V_rest_* and hyperpolarized *AP_thr_*. While such clustering analyses are limited by the somewhat low resolution data currently available, our approach is easily extensible to novel datasets, including from raw electrophysiological traces or additional data modalities like gene expression 36, 37 or morphology [5].

### Electrophysiological standards will improve future metaanalyses

A major goal of our project was to rigorously identify the sources of variance that limit the comparison of cross-study electrophysiological data. However, a difficulty that we regularly encountered came from the lack of formal standards used in reporting electrophysiological data. For example, during the NeuroElectro database QC audit (see Appendix), we observed at least six different definitions for calculating R_*input*_ (Fig. S4). Rigorously accounting for such inconsistent definitions was further hindered by frequently insufficient methodological details describing how each property was defined and calculated (i.e., only ∼ 60% of electrophysiological measurements were described with adequate detail to enable independent re-measurement). Thus, to the extent that inconsistent electrophysiological definitions yield systematically different measurements, naively pooling across studies (as we have done here) will continue to be a source of unexplained variance until more complete reporting standards are adopted.

Similarly, our approach requires mapping each extracted datum to a canonical neuron type. Since investigators use different terminologies to refer to neuron types [6], we used the community-generated expert-defined list of neuron types provided by NeuroLex 17, 18. This choice saves us from the challenging task of redefining the canonical list of neuron types, but at present these definitions currently “lump” rather than “split” neuron types (e.g., “neocortex layer 5-6 pyramidal neuron”). While this lumping will also add unexplained variance to neuron type biophysical measurements, we have built the mapping of data to neuron type in NeuroElectro to be highly flexible, allowing NeuroElectro to similarly evolve to match updates in neuron type definitions.

Based on our experiences, we recommend the usage of common definitions for basic biophysical measurements [13] and neuron types 6, 18. We also ask that experimentalists report more basic electrophysiological information within articles and, if possible, publish such data using machine-readable formats like data tables. Similarly, relevant experimental details should be clearly stated within methods sections [12] (e.g., liquid junction potential correction and recording quality criteria). In contrast to mandating that investigators standardize experimental protocols (e.g., using the same mouse line or electrode pipette solution), these shifts in data reporting practices we propose are simple, requiring minimal changes to current workflows. Implementing these basic recommendations will facilitate further data compilation efforts and the ultimate development of a comprehensive “parts list” of the brain [8].

### Meta-analysis as a remedy for the reproducibility crisis in neuroscience

Biomedical science is perceived to be undergoing a “reproducibility crisis” 11, 12, where up to half of published findings may be false [10]. In neurophysiology, such irreproducibility has been used to justify efforts by large single institutes to standardize the recollection of large amounts of data in the absence of an overarching question or hypothesis 7, 9, 38].

We feel that our meta-analysis approach offers a potential alternative solution. Specifically, by aggregating vast amounts of previously collected quantitive data and tagging these with appropriate experimental metadata, the metadata can help resolve systematic discrepancies between data values. Thus, as opposed to the standard practice of only utilizing data from a single study or laboratory, this “wisdom of the crowds” approach explicitly links together the work of a wide community of investigators. Neuropsychiatric genetics provides an excellent example, where investigators have identified greater numbers of genetic loci conferring significant disease risk by pooling subject data across sites and consortia [37]. While such quantitative meta-analyses are in their infancy in cellular and systems neuroscience 1, 2, we feel that this approach increases the reach and impact of any one publication and has the potential to greatly increase the rate of progress in our field.

## Materials and Methods

### Electrophysiological database construction

We built a custom infrastructure for extracting neuron type-specific electrophysiological measurements, such as *R_input_* and *AP_hw_,* as well as associated metadata (detailed in [14]). Briefly, our methods for obtaining this information are as follows. First, we obtained thousands of full article texts as HTML files from publisher websites. We next searched for articles containing structured HTML data tables; within these tables we used text-matching tools to find entities corresponding to electrophysiological concepts like “input resistance” and “spike half-width”. Our methodology accounts for common synonyms and abbreviations of these properties (e.g., “input resistance” is often abbreviated as *“ R_in_*”).

To identify measurements from neuron types collected in control or normotypic conditions, we used primarily manual curation. We used the listing of vertebrate neuron types provided by NeuroLex 17, 18 (http://neurolex.org/wiki/Vertebrate_Neuron_OVerview) to link mentions of a neuron type within an article to a canonical, expert-defined neuron type. After identifying both neuron type and electrophysiological concepts, we then extracted the mean electrophysiological measurement (and when possible, the error term and number of replicates). In most cases, however, our current methods were unable to extract the number of replicates; we thus have limited our focus here to analyses using mean measurements alone. Following application of the automated algorithms, we manually curated the extracted information and standardized electrophysiological property measurements to the same overall calculation methodology (e.g., Fig. S1C,D). In addition, we also used manual curation alone to extract information from ~35 articles which did not contain information in a formatted data table (typically older articles only available as PDFs or articles specific to olfactory circuit physiology).

We also obtained information on article-specific experimental conditions from each article's methods section. Specifically, we considered the effect of: animal species, animal strain (here we distinguished between strains of rats but not different genetic strains of mice), electrode type (sharp versus patch clamp), preparation type *(in vitro, in vivo,* cell culture), liquid junction potential correction (explicitly corrected, explicitly not corrected, not reported in manuscript), animal age (in days; when only animal weight was reported, we manually converted reported weight to an approximate age using conversion tables provided by animal vendors), and recording temperature (we assigned reports of “room temperature” recordings to 22°C and in *vivo* recordings to 37 C). Additional methodological details, including recording and electrode solution contents and pipette resistances, will be considered in future iterations.

Explicit instructions and details for the quality control (QC) audit of the NeuroElectro database are provided in the Appendix.

### Data analysis

#### Data filtering and preprocessing

Before performing systematic analyses of the data within the NeuroElectro database (i.e., data referred to following Fig. 2 onwards, unless otherwise specified), we performed the following filtering steps: 1) we excluded non-brain neuron types (e.g., we excluded spinal cord neuron types); 2) we excluded data collected from dissociated and slice cell cultures; 3) to account for large differences in animal age across species, we only used data from rats, mice, and guinea pigs; 4) we excluded data from embryonic and perinatal (<5 days) animals; 5) due to inconsistencies in the definition of *τ_m_*, we excluded measurements of *τ_m_* which deviated greater than 2 standard deviations from the median measurement across articles. Where metadata attributes were not reported or were unidentifiable within an article (which was typically rare for the experimental conditions that we focused on), we used mean (or mode) imputation for continuous (or categorical) metadata attributes [39].

#### Metadata incorporation

We used statistical models to account for the influence between experimental metadata and measured electrophysiological values. Specifically, we modeled the relationship between electrophysiological measurements and experimental metadata as 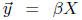 where 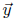 denotes the vector of electrophysiological measurements corresponding to a single property across all articles (e.g., V_*rest*_); *X* denotes the regressor matrix where rows denote the metadata attributes associated with a single measurement *yi* (e.g., 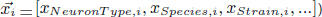 and *β* are the regression coefficients denoting the relative contribution of each metadata attribute. We logio-transformed measurements of *R_input_, *τ*_*m*_, AP_*hw*_*, and animal age to normalize values because these varied across multiple orders of magnitude and/or to enforce that these values remain strictly positive following metadata-based adjustment.

When combining the influence of multiple metadata attributes into a single regression model (Fig. 2C), we wished to use powerful and flexible models to capture the relationship between metadata and measurement variance while also mitigating the tendency of more complex statistical models to overfit the data. Thus, when fitting statistical models, we used stepwise regression methods (implemented as LinearModel.stepwise in MATLAB) to add model terms one-by-one and added terms until the model's Bayesian Information Criterion (BIC) was optimized. Our choice of BIC here is based on its conservativeness relative to other approaches for model selection, which helps protect against statistical overfitting. Furthermore, for each electrophysiological property, we selected the potential model complexity from a set of candidate models (i.e., models that included terms for only: constant, linear, purely quadratic, interaction, interaction + quadratic). We selected model complexity using 10-fold cross-validation and minimization of the sum of squared errors on out-of-sample data. In reporting the variance explained by different models, we used adj. R^2^ to compare between models differing in their number of parameters.

After fitting metadata regression models for each electrophysiological property, we adjusted each electrophysiological measurement to its estimated value had it been measured under conditions described by the population mean metadata value (or mode for categorical metadata attributes). For example, since the majority of data were recorded using patch clamp electrodes, we then adjusted measurements made using sharp electrodes to their predicted value had they been recorded using patch clamp electrodes. To assess the robustness of the fit of the regression models, we re-ran the regression analysis on different versions of the dataset where the data were randomly subsampled (Fig. S8). Note that the penalty that BIC imposes against overfitting is stronger when there are fewer data points used to fit the models. Thus, progressively subsampling the dataset penalizes away the amount of variance in the electrophysiological data that can be explained by experimental metadata.

#### Electrophysiological property correlation and neuron type similarity analyses

For analysis of electrophysiological and neuron type correlations, we first pooled data by averaging measurements collected within the same neuron type. We then defined each neuron type using its vector of six electrophysiological measurements. We quantified correlations between pairs of electrophysiological properties using Spearman's correlation, which assesses the rank-correlation and allows for detection of relationships that are monotonic but not necessarily linear. We used the Benjamini-Hochberg false discovery rate procedure to control for multiple comparisons performed in the pairwise correlation analysis.

To quantify how much variance across electrophysiological properties could be explained by subsequent principal components (PCs), we needed to first account for missing or unobserved measurements within our dataset. For example, some neurons did not have a reported measurement for *τ_m_* or *AP_thr_* within our dataset. To address this issue of missing data [39], we used pPCA, a modification of traditional PCA that is robust to missing data. To further mitigate the problem of missing data, in this analysis we only considered neuron types that were defined by at least three different articles and with no more than two of the six total electrophysiological properties missing; after this filtering step, less than 10% of total electrophysiological observations were missing.

To quantify the electrophysiological similarity of neuron types, we calculated the pairwise Euclidean distances between pairs of neuron types defined by the vector of six electrophysiological properties and used a dendrogram analysis to sort neuron types on the basis of electrophysiological similarity. Missing or unobserved electrophysiological measurements were imputed using pPCA, as described above. Here, we chose to be agnostic about the relative importance of each biophysical property and weighted biophysical properties based solely on their relative measurement uncertainty (defined in Fig. 2C). Thus, properties which tend to show greater cross-study variability (such as *τ_m_* will be less down weighted in this analysis relative to more reliable measurements like *V_rest_.* Empirically, we found this weighting to help further mitigate unaccounted-for measurement and methodological variability.

The dendrogram, *D,* denoting the hierarchical similarity among neuron types, was constructed using linkages computed by Ward's minimum variance method. We used multiscale bootstrap resampling to assess the statistical significance of subtrees of *D* using the pvclust package in the language R 40, 41 (referred to as the AU p-value in Fig. 5). A detailed description of the pvclust algorithm methodology is provided in the Appendix.

### Acute slice electrophysiology

#### Animals

Hippocampal CA1 pyramidal cell recordings were conducted using postnatal day (P)15-18 M72-GFP mice [42] and Thy1-YFP-G mice [43]. Main olfactory bulb mitral cell recordings were conducted using P15-18 M72-GFP, Thy1-YFP-G, and C57BL/6 mice. A subset of data from these neurons has been published previously [44]. Main olfactory bulb granule cell recordings were conducted using P18-22 C57BL/6 and albino C57BL/6 mice. Neocortical basket cell recordings were conducted using a P26 parvalbumin reporter mouse, resulting from a cross between Pvalb-2A-Cre (Allen Institute for Brain Science) and Ai3 [45] lines. Striatal medium spiny neuron recordings were conducted using P14-17 M72-GFP mice. A total of 20 mice of both sexes were used in this study. Animals were housed with litter mates in a 12/12 hr light/dark cycle. All experiments were completed in compliance with the guidelines established by the Institutional Animal Care and Use Committee of Carnegie Mellon University.

#### Slice preparation

Mice were anesthetized with isoflurane and decapitated into ice-cold oxygenated dissection solution containing (in mM): 125 NaCl, 25 glucose, 2.5 KCl, 25 NaHCO3, 1.25 NaH2PO4, 3 MgCl2, 1 CaCl2. Brains were rapidly isolated and acute slices (310 Am thick) prepared using a vibratome (VT1200S; Leica or 5000mz-2; Campden). Slices recovered for 30 min in ∼37°C oxygenated Ringer's solution that was identical to the dissection solution except for lower Mg concentrations (1 mM MgCl2) and higher Ca^2+^ concentrations (2 mM CaCl2). Slices were then stored in room temperature oxygenated Ringer's solution until recording. Parasagittal slices were used for hippocampal and striatal recordings. Coronal slices were used for neocortical recordings. Horizontal slices were used for main olfactory bulb recordings.

#### Recording

Slices were continuously superfused with oxygenated Ringer solution warmed to 37°C during recording. Cells were visualized using infrared differential interference contrast video microscopy. Hippocampal CA1 pyramidal cells (n=10) and were identified by their large soma size, pyramidal shape, and position within CA1. Neocortical basket cells (n=5) were identified by expression of YFP fluorescence. Main olfactory bulb mitral cells (n=10) were identified by their large cell body size and position within the mitral cell layer. Main olfactory bulb granule cells (n=9) were identified by their small cell body size and position within the mitral cell or granule cell layers. Striatal medium spiny neurons (n=8) were identified by their extensively spine-studded dendritic arbors viewable under epifluorescence through Alexa Fluor 594 cell fills. Whole-cell patch clamp recordings were made using electrodes (final electrode resistance: 6.1±1.1 MΩ, μ ± σ; range: 4.4-8.7 MΩ) filled with (in mM): 120 K-gluconate, 2 KCl, 10 HEPES, 10 Na-phosphocreatine, 4 Mg-ATP, 0.3 Na3GTP, 0.2-1.0 EGTA, 0-0.25 Alexa Fluor 594 (Life Technologies), and 0.2% Neurobiotin (Vector Labs). Cell morphology was reconstructed under a 100X oil-immersion objective with Neurolucida (MBF Bioscience). No cells included in this dataset exhibited gross morphological truncations. Mitral cells were recorded in the presence of CNQX (10 *μM),* DL-APV (50 *μM),* and Gabazine (10 μM) to limit the influence of spontaneous synaptic long-lasting depolarizations [46] on measurements of biophysical properties. Data were low-pass filtered at 4 kHz and digitized as 10 kHz using a MultiClamp 700A amplifier (Molecular Devices) and an ITC-18 acquisition board (Instrutech) controlled by custom software written in IGOR Pro (WaveMetrics). The liquid junction potential (12-14 mV) was not corrected for. Pipette capacitance was neutralized and series resistance was compensated using the MultiClamp Bridge

Balance operation and frequently checked for stability during recordings. Series resistance was maintained below ∼20 MΩ for all pyramidal cell, mitral cell, basket cell, and medium spiny neuron recordings (14.0±2.7 MΩ, *μ* ± *σ;* range: 8.4-20.1 MΩ). Higher series resistance (30.7±6.2 MΩ, μ ± σ range: 23.5-43.0 MΩ) was permitted in granule cell recordings due to their small (∼10 soma sizes. After determination of each cell's native *V_rest_,* current was injected to normalize V*_rest_* to — 65, —58, —65, —70, and —80 mV for pyramidal cells, mitral cells, granule cells, basket cells, and medium spiny neurons, respectively, before determination of other biophysical properties. In preliminary experiments, we also recorded from layer 5/6 neocortical pyramidal cells. However, these recordings were not further pursued due to the extensive electrophysiological and morphological heterogeneity observed within this broad category of neuron type.

#### Analysis

*V_rest_* was determined immediately after cell break in. *τ_m_* was calculated from a single-exponential fit to the initial membrane potential response to a hyperpolarizing step current injection. *R_input_* was calculated as the slope of the relationship between a series of hyperpolarizing step current amplitudes (that evoked negligible membrane potential “sag”) and the steady-state response of the membrane potential to injections of those step currents. In a subset of recordings, *R_input_* was also calculated as the steady state response of the membrane potential to a single step current injection (evoking a ∼5 mV hyperpolarization) averaged across 50 trials. Both methods yielded equivalent results. To determine action potential properties of each neuron, a series of 2 s-long depolarizing step currents was injected into the neuron. The first action potential evoked by the weakest suprathreshold step current (i.e., the rheobase input) was used to determine the action potential properties of the neuron. *AP_thr_* was calculated as the first point where the membrane potential derivative exceeded 20 mV/ms. *AP_amp_* was measured from the point of threshold crossing to the peak voltage reached during the action potential. This amplitude was then used to determine *AP_hw_*, calculated as the full action potential width at half maximum amplitude of the action potential.

For the confusion matrix analysis (Fig. 3), we used a Euclidean distance approach identical to that used in the analysis of electrophysiological neuron type similarity. Specifically, we represented each recorded single-cell via its measurements along the six major electrophysiological properties. We then compared the similarity of each recorded neuron to the mean electrophysiological measurements of each of the five corresponding “canonical” neuron types from NeuroElectro, after either: first adjusting the filtered NeuroElectro data to the methodological conditions used in our laboratory; or using the raw and unfiltered data values from NeuroElectro, unnormalized for methodological differences. We classified each recorded single-cell to the most similar of the canonical NeuroElectro neuron types by finding the NeuroElectro neuron type with the smallest Euclidean distance to the single-cell, forming the basis for the confusion matrix.

## Data and code availability

The NeuroElectro dataset and spreadsheets listing mined publications, neuron types, electrophysiological properties are provided at http://neuroelectro.org/static/src/ad/paper/supp/material.zip. The analysis code used here is available at http://github.com/neuroelectro.

## ACKNOWLEDGMENTS

We thank R. Kass, A-M. Oswald, A. Gittis, A. Barth, E. Fanselow, and P. Pavlidis for helpful discussions and comments on the manuscript. We additionally thank A. Gittis for guidance on targeted neocortical basket cell recordings; G. LaRocca for technical support; and D. Tebaykin for NeuroElectro web-development assistance. We are especially grateful to all of the investigators whose collected data are represented within the NeuroElectro database. This work was supported by a NSF Graduate Fellowship (to S.J.T.), NIDCD award F31DC013490 (to S.D.B.), NIDCD award F32DC010535 and NIMH award R01MH081905 (to R.C.G.), and NIDCD Grant R01DC005798 (to N.N.U.).

## Appendix

### Electrophysiological database quality control assessment

To validate and quality control (QC) the accuracy of our semiautomated data extraction methods, we conducted a systematic audit of a randomly chosen 10% of the algorithmically-mined articles (n = 27 articles) which had curated electrophysiological data obtained from a structured data table within an article.

Specifically, four curators (two pairs of two curators, each working independently) were each tasked with validating the accuracy of concept identification, data extraction, and data standardization. The curators (S.J.T., S.D.B., M.G., and R.C.G.), each had extensive experience reading literature and designing and performing electrophysiology experiments. Curators were split into pairs of two curators each, where each member of each pair independently curated the same manuscripts, allowing an assessment of inter-curator consistency.

Below are listed the explicit instructions provided to each curator during the 10% article QC audit to independently validate the accuracy of the NeuroElectro database.

1. Assess whether each neuron type mentioned within a HTML data table had been mapped correctly (“yes”, “no”, or ‘'ambiguous”), based on the provided listing of canonical neuron types and their definitions. “Ambiguous” responses and notations were used in cases where the neuron type was not included in the provided neuron type list.
2. Assess whether electrophysiological property concepts had been identified and mapped correctly (“yes”, “no”, or “ambiguous”). In addition, curators were asked to explicitly describe how each electrophysiological property had been calculated, and evaluate whether the electrophysiological measurement could be in principle replicated based on the provided methodological details. For example, for AP_*thr*_, if the article defined how threshold was computed (e.g., voltage derivative threshold criterion) but did not describe how spikes were evoked (e.g., rheobase current injection), then the AP_*thr*_ measurement was judged as non-replicable. Curators were instructed to attempt to find such electrophysiological definitions if they were referenced within a previous manuscript. Curators were not allowed to report “ambiguous” for evaluations of replicability.
3. Extract the electrophysiological data corresponding to each neuron type-electrophysiological property pair as it directly appears in the formatted data table. Following this extraction, standardize the data value to the common calculation methodology (e.g, measurements of input resistance reported in GO were standardized to MO).
4. Assess whether methodological metadata concepts were identified and mapped correctly. In addition, curators were asked to curate information for a small number of additional metadata concepts (extracellular *Mg^2+^* and *Ca^2+^* concentrations, slice thickness, etc.) as seed data for future metadata extraction algorithms.

To evaluate the outcome of the QC audit, we quantified agreement between curators and between curators and NeuroElectro using a simple percentage agreement measure. Concepts which were labelled as ambiguous by the curator were not considered in the quantification. Because we quantify NeuroElectro accuracy in comparison to the human curators, the inter-curator agreement measure sets a rough upper bound on the potential maximum accuracy of NeuroElectro. Specifically, the aggregate intercurator consistency measure of 95% sets the upper bound of NeuroElectro accuracy at 97.5%.

*QC results.* We found close agreement between the manually curated QC data and NeuroElectro, with 96% accuracy of the NeuroElectro database overall (specifically: 93% for neuron types, 99% for electrophysiological concepts, 96% for experimental conditions, and 95% for correctly extracted and standardized electrophysiological data). Inter-curator agreement was also high, with 95% of concepts and data identified and extracted identically across each pair of two curators. Moreover, “mistakes” or miscurated entries within the NeuroElectro database usually represented cases where the underlying concept or data were truly ambiguous (e.g., a neuron type which did not explicitly exist within the NeuroLex list of neuron types).

Within the QC sample, we also analyzed how often authors use different definitions for similar electrophysiological properties and whether sufficient details were provided to independently replicate each measurement. Strikingly, we found that sufficient methodological details were provided to replicate only 42% of reported electrophysiological measurements. While this measure of replicability is inherently subjective and dependent on electrophysiological experience (yielding an inter-curator agreement of only 65%), our results nevertheless parallel other aggregate measures of methodological rigor, such as antibody reporting [12].

### Description of dendrogram bootstrap resampling

We used multiscale bootstrap resampling to assess the statistical significance of subtrees of *D* using the pvclust package in the language R[40]. The pvclust dendrogram multiscale bootstrap resampling algorithm proceeds as follows: specifically, given an n *x* p data matrix M (here, n refers to neuron types and *p* refers to the six electrophysiological properties), pvclust first generates a number of bootstrapped versions of *M* through randomly sampling columns from M with replacement (10,000 bootstrap samples were used). For each bootstrapped data matrix, *Mi*, a dendrogram D_*i*_ was generated through hierarchical clustering. Next, for each subtree in the original dendrogram D, the analysis assesses how often the same subtree appears across the bootstrapped dendrograms D_1:10_,_000_. Here, subtree equality is defined by subtrees that share identical tree topology and neuron membership but does not assess equality of branch lengths. Lastly, because the bootstrap probability is known to be a downwardly biased measure for determining subtree probability [41], pvclust corrects for this downward bias by performing the entire bootstrap procedure multiple times at a number of scales by resampling M to have differing numbers of columns (here, we use 3 through 9 columns in M). This allows for the bootstrap probability to be corrected, yielding the approximately unbiased p-value for each subtree.

**Supp. Fig. 1.**
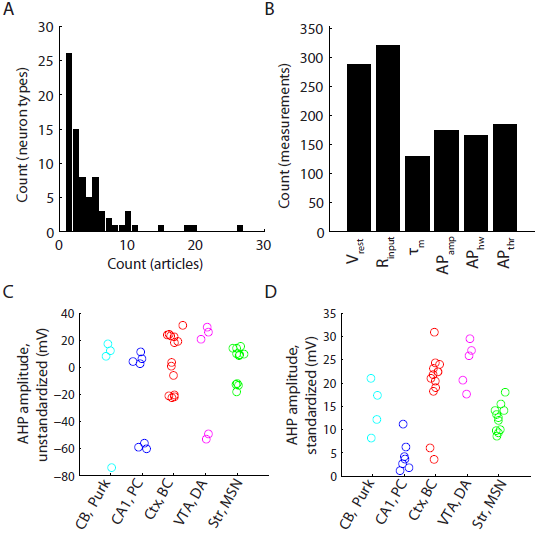
Distribution of neuron types and electrophysiological properties represented in NeuroElectro and illustration of electrophysiological property standardization. A) Frequency histogram of distribution of neuron types versus number of articles containing information about each neuron type. B) Count of unique measurements of the six electrophysiological properties explored in this manuscript. C,D) Illustration of manual electrophysiological property standardization for NeuroElectro measurements extracted from literature. Example afterhyperpolarization (AHP) amplitude measurements before (C) and after standardization (D) to a common calculation definition. Neurons plotted are cerebellar Purkinje cells, CA1 pyramidal cells, cortical basket cells, ventral tegmental area dopaminergic cells, and striatal medium spiny neurons (abbreviated as Purk; CA1, pyr; Ctx, bskt; VTA, DA; and Str, MSN; respectively). Each circle denotes the value of the mean electrophysiological measurement reported within an article.

**Supp. Fig. 2.**
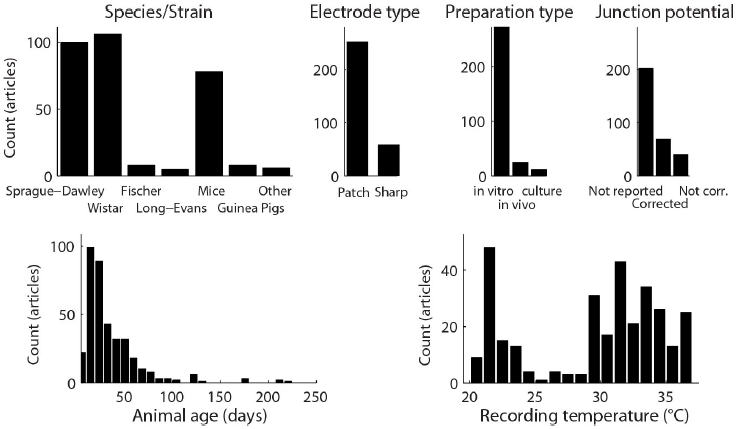

**Supp. Fig. 3.**
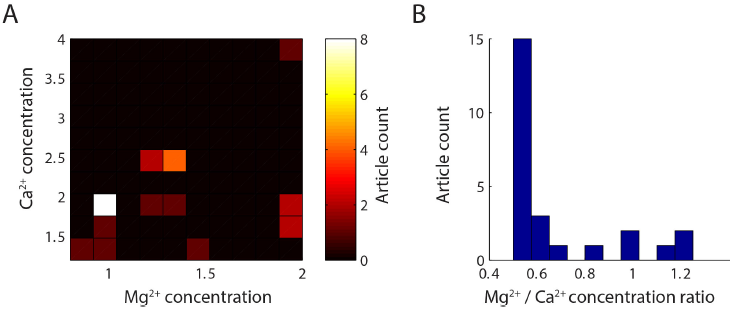
*Quantification of Mg^2+^ and* Ca^2+^ *recording solution concentrations among articles in quality control subset.* A) *2*dimensional histogram of Mg^2+^ and Ca^2+^ recording solution concentrations, reported in mM. The most commonly reported concentration pair is 1mM Mg^2+^ and 2 mM Ca^2^+. B) Same as A, but reported as ratio of Mg^2+^ to Ca^2+^ concentration. n = 27 articles quantified; 2 articles not shown since *in vivo* recording conditions were used and no external recording solution was reported.

**Supp. Fig. 4.**
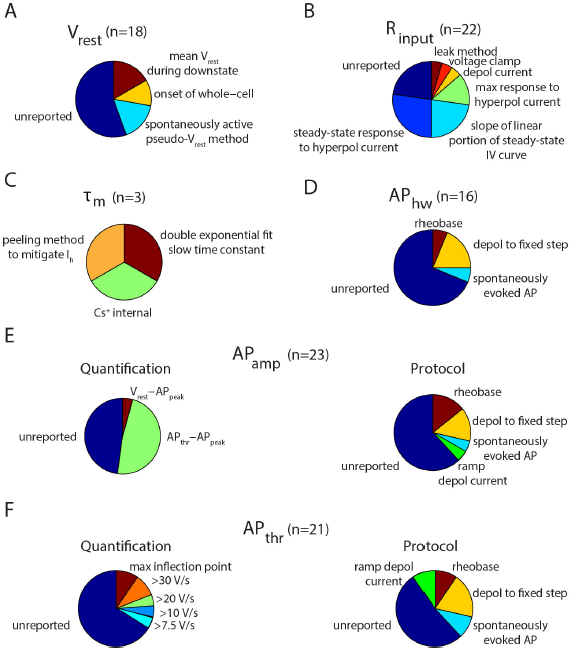
*Compilation of different overall methods for calculating electrophysiological properties from the sample of curated articles in the QC audit.* A-F) Pie charts and labels indicate breakdown of electrophysiological calculation methodology and n indicates number of property measurements found in sample. Label ‘unreported’ indicates that no specific methodological description could be found. n = 27 articles quantified in QC subset. A) Resting membrane potential *(V_rest_),* label ‘spontaneously active pseudo-*V_rest_* method’ indicates methodology for quantifying *V_rest_* in spontaneously active neurons. B) Input resistance *(R_input_*), label ‘leak method’ indicates method for calculating *Ri_n_p_u_t* based on leak current. C) Membrane time constant (τ_*m*_), label ‘peeling method to mitigate *Ih* ’ indicates method calculating *τ_m_* that corrects for sag current influence, label ‘*Cs+* indicates the use of cesium ions in the electrode pipette solution.; D) Action potential half width *(AP_hw_*). Labels indicate different protocols for eliciting spikes from which *AP_hw_* is calculated. By definition, all *AP_hw_* measurements have been quantified as AP full width at half maximal amplitude, usually from the first evoked AP in train. E) Action potential amplitude *(AP_amp_*). Pie charts indicate methodology for quantifying *AP_amp_* (left) or protocol used to elicit action potentials (right). Quantification labels indicate whether *AP_amp_* is defined as the difference between AP threshold and peak or *V_rest_* and AP peak. F) Action potential threshold *(AP_thr_*), label ‘max inflection point’ indicates identification of action potential threshold via *2^nd^* derivative of voltage.

**Supp. Fig. 5.**
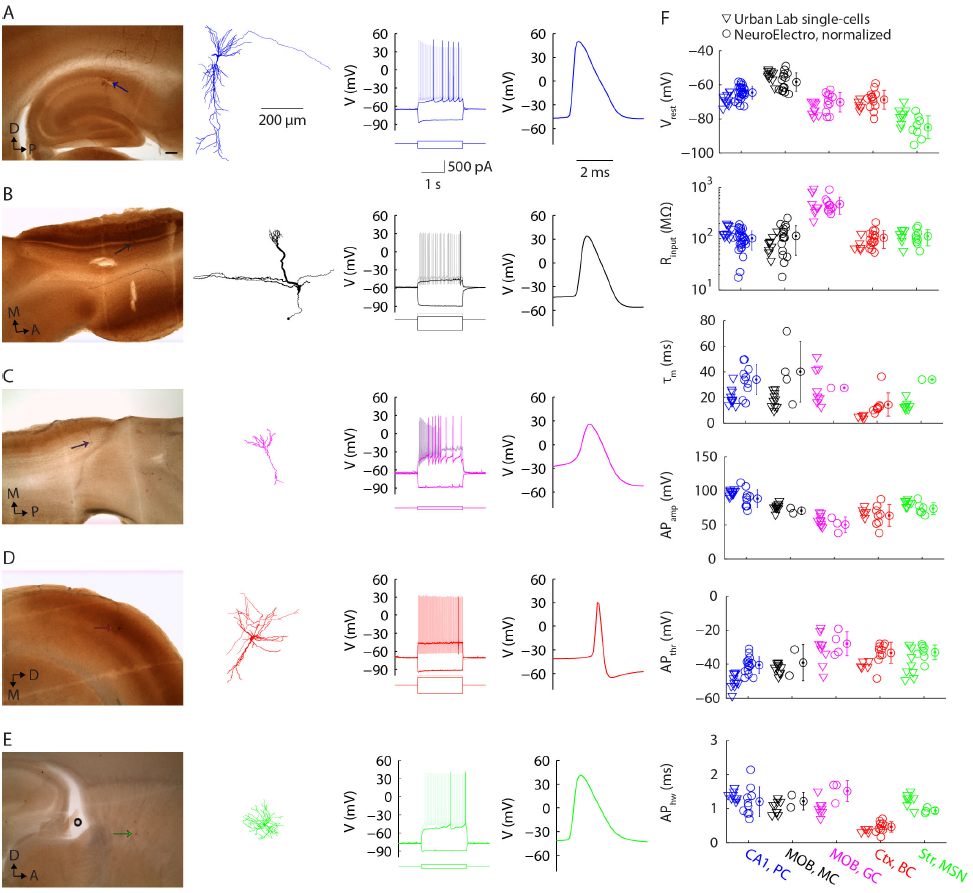
*Validation of NeuroElectro database measurements with collection of raw data.* A) Representative targeted recording of a hippocampal CA1 pyramidal cell (“CA1, PC”), showing anatomical position and morphological reconstruction (left), response to hyperpolarizing and depolarizing rheobase and suprathreshold step current injections (middle), and action potential waveform (right). Anatomical scalebar: 200 *μ*m. B-D) Same as A for: main olfactory bulb mitral cell (B; “MOB, MC”), main olfactory bulb granule cell (C; “MOB, GC”), neocortical basket cell (D; “Ctx, BC”), and striatal medium spiny neuron (E; “Str, MSN”). F) Summary of targeted *in vitro* recordings and comparison to text-mined, metadata-adjusted values from NeuroElectro. Abbreviations: dorsal (D), posterior (P), medial (M), anterior (A). Morphological reconstructions (except the representative granule cell) have been moderately thickened to aid visualization of thinner

**Supp. Fig. 6.**
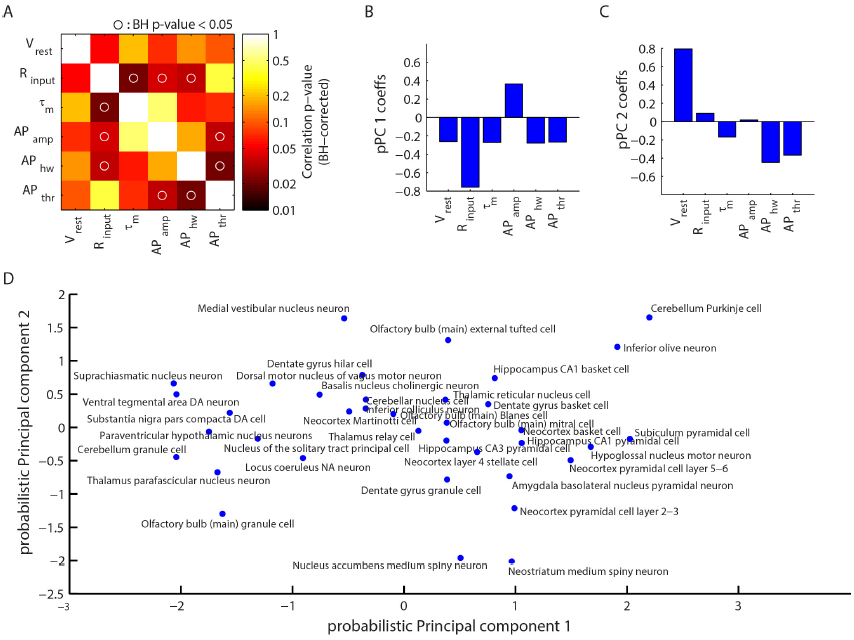
*Expanded analysis of correlations among electrophysiological properties.* A) Benjamini-Hochberg adjusted P-values for pairwise electrophysiological property correlation matrix shown in Fig. 4. B,C) Coefficients corresponding to the first (B) and second (C) probabilistic principal component (pPC). D) Projection of neuron types onto space defined by first and second pPCs. Note that the first pPC qualitatively reflects the axis of electrotonically small (left) vs. large (right) neuron types, while the second pPC qualitatively reflects the axis of basal excitability of neuron types, separating hyperpolarized (bottom) from depolarized (top) resting membrane potentials.

**Supp. Fig. 7.**
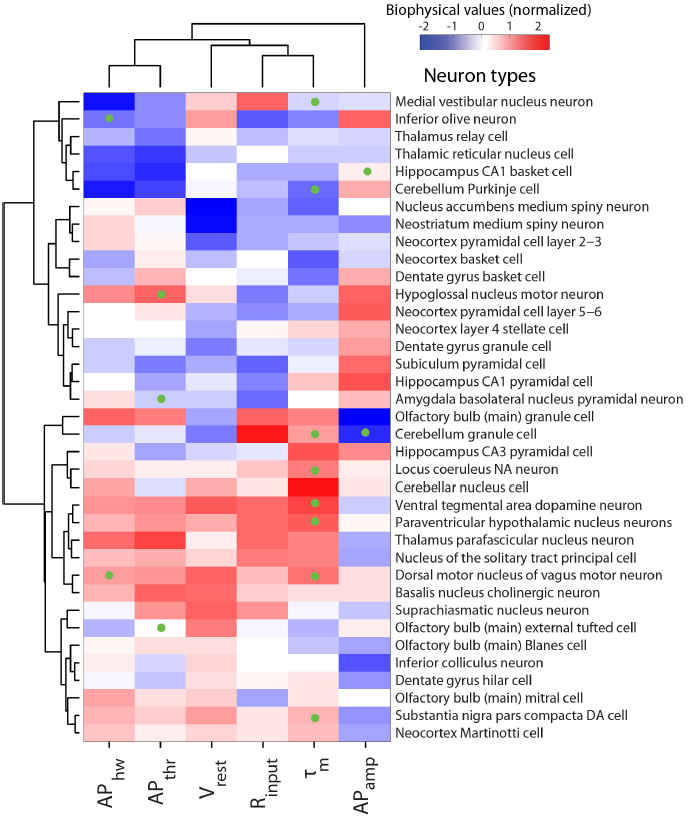
*Hierarchical clustering of neuron types, without first normalizing for differences in experimental metadata.* Same as Fig. 5, but computed for biophysical data without first adjusting for differences in experimental conditions. Neuron types sorted in order of biophysical similarity (similiarity indicated by levels of dendrogram; dendrogram linkages computed using Ward's method and Euclidean distances). Heatmap values indicate observed neuron type-specific electrophysiological measurements, red (blue) values indicate large (small) values relative to mean across neuron types. Only neuron types with measurements defined by at least three articles and with at least four (of the six total) biophysical properties reported were used in this analysis. Probabalistic PCA was used to impute unobserved measurements, indicated via green dots on heatmap.

**Supp. Fig. 8.**
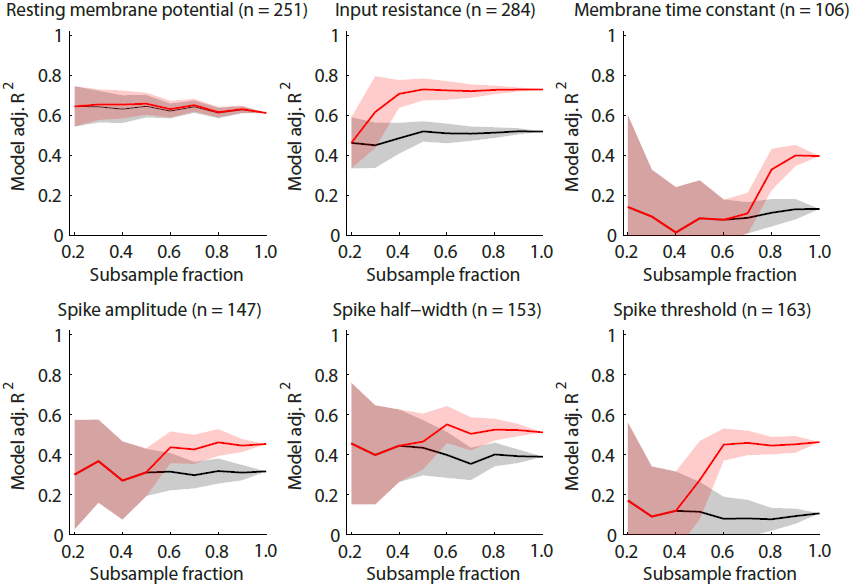
*Influence of dataset size versus predictive power of metadata for explaining electrophysiological measurement variability.* Panels show different electrophysiological properties and lines show explanatory power of statistical model when using neuron type information only (black) or neuron type plus all metadata (red) as a function of randomly subsampling the original dataset to smaller sizes (on abscissa). Original dataset size indicated by subsample fraction = 1.0. Shaded lines indicate standard deviations when resampling the dataset 25 times per subsampling size. Note that as dataset is subsampled to smaller sizes, explanatory power of the model that includes metadata is not greater than the model which includes neuron type information only.

